# B-cell specific expression of the murine *Myd88^L252P^* mutation correlates with the establishment of an immunosuppressive microenvironnement in Waldenström macroglobulinemia lymphoplasmacytic lymphoma

**DOI:** 10.1101/2023.09.28.559899

**Authors:** Quentin Lemasson, Ophélie Téteau, Mina Chabaud, Laurie Jeanson, Jean Feuillard, Christelle Vincent-Fabert

## Abstract

Waldenström macroglobulinemia is a rare and indolent lymphoproliferative disorder genetically characterized by the presence of the L265P mutation in the *MYD88* gene in nearly each case. Despite its slow progression, Waldenström macroglobulinemia remain incurable due to the lack of specific treatments. The importance of the tumour microenvironment was well documented since the last decade especially concerning the study of solid tumours, but the failures of the immune microenvironment are also experiencing growing interest for B cell lymphomas. In this study, we investigated the implication of some dysregulations of the immune microenvironment in Waldenström macroglobulinemia through our *Myd88^L252P^* mutated transgenic model, which may explain its progression. In essence, we highlighted the existence of multiple immune escape mechanisms that lead to T-cell exhaustion and participate to Waldenström disease progression. This work in animals opens up new prospects for the development of new therapeutic combinations in WM targeting reactivation of the immune system.

## Introduction

Waldenström’s disease (WM), described by Jan G. Waldenström in 1944, is an incurable, indolent lymphoplasmacytic lymphoma (LPL) of the elderly that accounts for less than 5% of B-cell lymphomas. This slowly evolving lymphoma is characterized by expansion of a lymphoplasmacytic clone of medullary localization with secretion of a monoclonal immunoglobulin type M (IgM). Being specific of WM, clonal cells exhibit a continuous differentiation between the small mature lymphocyte and the mature plasma cell (PC). We recently demonstrated that this unique characteristic may be used as a biological marker for WM diagnosis by evidencing the B-cell clonality in both lymphocytes and plasma cell by flow cytometry [1]. Similar to other lymphomas, cytogenetic and molecular abnormalities have been identified and can be used as prognostic factors. Indeed, the main chromosomal aberrations are 6q deletions (in 20-40% of patients), 13q deletions (10-15%), trisomy 18 (10%), trisomy 4 and 17p deletions (8%) [2]. Using Next-Generation Sequencing (NGS), the discovery of the L265P mutation in *MYD88* (an adaptor protein involved in Toll Like Receptor signalling and activating the NF-kappa B (NF-κB) pathway) has demonstrated that WM is a true entity genetically distinct from other indolent lymphomas with IgM secretion [3]–[5]. Indeed, WM is associated with an activating mutation of *MYD88* in more than 90% of cases (*MYD88 L265P*, the most frequent). Another frequent mutation (30% of WM patients), almost specific of WM, is the WHIM-like mutation of CXCR4, which results in a lack of desensitisation of this receptor, which would explain why WM B cells reside in the bone marrow [6].

Despite current progress and the development of new therapies, WM remains still incurable [7]. It is therefore important to develop new approaches in this field. Until recently, adequate preclinical models, which is an important step to develop these new approaches were lacking. With few other groups, we have created an animal model which intends to mimic WM [8]–[11]. Our strategy consisted in inserting the *Myd88^L252P^* mutation (mouse orthologue of the human mutation) in the *Rosa26* locus. These mice were then crossed with *Cd19^Cre^* mice to induce expression of the mutation only in B cells using the Cre/LoxP system (*Myd88^L252P^* model, respecting the endogenous mouse MYD88 signalling and forcing expression of the mutant in B cells in a pseudo-heterozygote manner) [11]. Our *Myd88^L252P^* mice developed at 9-13 months of age (i) an indolent, (ii) lymphoplasmacytic B-cell lymphoma in the spleen (iii) with, in the blood, secretion of a monoclonal IgM and (iv) a lymphoplasmocytic B-cell expansion in the bone marrow. For the first time, we have created a mouse model that recapitulates the main features of WM [11]. This is a unique opportunity to both understand how MYD88 induces WM and to identify new therapeutic perspectives.

Escape from the immune response is a crucial step in the emergence of lymphomas, especially for those with NF-κB activation [12]–[15]. For this reason, tumour escape has been a popular area of study over the past decade, enabling the emergence of innovative therapies whose aim is no longer to target the tumour directly, but to reactivate the immune system. Indeed, in order to survive and proliferate, cancer cells often reprogram the tumour microenvironment (TME) to protect themselves from the defense mechanisms of the host immune system. Alterations in direct cell-cell interactions or cytokine production have been reported in many cancers, including lymphomas [16]. Among the mechanisms of escape from the immune response, LT exhaustion through overexpression of immune checkpoints and/or secretion of immunosuppressive cytokines have been reported in some indolent lymphomas [17]–[21]. However, when it comes to WM, the role of the immune microenvironment in its development remains poorly understood. Knowing the pathways deregulated in WM cases (TLR/NF-κB, JAK/STAT, BCR and PI3K/AKT signalling) and their consequences in terms of cytokines secretion, the tumour microenvironnement should be disturbed. Moroever, the indolent form of this disorder suggests that WM cells should develop many mechanisms for shutting down the immune system. In 2013, Grote D. et al. showed for the first time that PD-L1 and PD-L2 were overexpressed not only on the surface of WM tumour cells, but also on the surface of cells in the tumour microenvironment (dendritic cells, monocytes and macrophages), promoting their proliferation [22]. In 2018, Jalali S. et al. showed in patients that CD19^+^CD138^+^ WM cells overexpressed PD-1 ligands (PD-L1 and PD-L2) and, in addition, secreted large quantities of them, which were found in blood and bone marrow. Moreover, this work showed that lymphocytes cultured in the presence of serum from WM patients had their proliferation severely compromised [23]. Finally, in 2021 and 2022, two works presented results concerning the immune microenvironment in WM. In the first, Kaushal A. et al. combined mass cytometry with the CITE-seq technique to demonstrate that a tumour clone-specific T immune response was established very early in the MGUS phase, and that throughout disease development T lymphocytes will progressively acquire an exhausted phenotype with loss of the naive T compartment and desensitization to IFNγ [24]. A little later, Sun H. et al. performed a high-throughput study of the WM microenvironment by analysing bone marrow samples in single cell RNAseq, and demonstrated a loss of NK cell cytotoxic activity [25]. In addition, he observed the presence of depleted CD4 and CD8 expressing T cells with loss of the naive compartment and increase in Regulatory T cells (Tregs) [25]. These recent studies demonstrate that immune escape seems to take place during the development of WM, leading to T-cell exhaustion and inactivity. While these studies are promising in terms of their therapeutic potential, they remain to be confirmed, and they raise the question of the exact mechanisms involved. In that view and based on our results on the mouse model *Myd88^L252P^* which developed WM-like tumours, we have started to study the immune microenvironment in tumour spleen. Our results showed the presence of several immune escape mechanisms in our mouse model of *Myd88* mutated lymphoplasmacytic lymphoma. These new data strongly suggest that the immune microenvironment plays an important role in the development of the disease but that it could also be one of the actors of their transformation, opening up perspectives for therapeutics trials targeting a reactivation of the immune system.

## Material and methods

### Mouse model

The mouse model used in this work is the *Myd88^L252P^-Cd19^Cre^* model developed by the laboratory and previously described (Ouk et al., 2021). For B-cell expression of the transgene, *Myd88^L252P^ ^tg/tg^* animals (C57Bl6/J genetic background) were crossed with *Cd19^Cre^ ^tg/tg^* mice (Balb/c background). The animals used in this study are *Myd88^L252P^ ^tg/+^ - Cd19^Cre^ ^tg/+^.* These mice will hereafter be referred to as *Myd88^L252P^* mice. Animals were housed at 21–23°C with a 12-hour light/dark cycle. All procedures were conducted under an approved protocol according to European guidelines for animal experimentation (French national authorization number: 8708503 and French ethics committee registration number APAFIS#14581-2018041009469362 v3).

### Flow cytometry

Spleen cells from *Cd19^Cre^* and *Myd88^L252P^* were filtered through a sterile nylon membrane. Cell suspensions were resuspended at 4°C in a labelling buffer (PBS, 1% BSA, 2mM EDTA) and labelled with fluorescent conjugated monoclonal antibodies listed in Supplementary Materials and Methods. The "Mouse Tregs detection kit" (Miltenyi Biotec®, Bergisch Gladbach, Germany) was used for regulatory T-cell staining. Membrane labeling (CD4 2µL + CD25 2µL, 20 min at 4°C) was performed on 1 million cells. Cells were then fixed-permeabilized (30 min at 4°C) before intracellular FoxP3 labeling (30 min at 4°C).

Labelled cells were analysed using a Cytoflex LX (Beckman Coulter®, Brea, CA). Results were analysed using Kaluza Flow Cytometry software 2.1 (Beckman Coulter®, Brea, CA).

### Immunochemistry

Paraffin embedded tissue sections (5µm) were carried out on the histology platform of the Biscem collaborative services unit at the University of Limoges (Inserm 042, CNRS 2015, Hospital University Center of Limoges) and were deparaffinized as follows: slides were immersed successively in xylene twice for 3 min, 3 times for 3 min in 100% ethanol, once for 3 min in 95% ethanol and 3 times in PBS for 5 min. Then, slides were immersed in citrate buffer pH7 and heated 4 times for 5 min 40 sec in a microwave at 800W. For T-cell staining a primary antibody against CD3 (SP7, Abcam®, Cambridge, UK) was used. Primary antibody revelation was performed with Zytochem Plus (HRP) Anti-Rabbit (DAB) kit (Zytomed systems®, Berlin, Germany) following supplier instructions. Image acquisition was performed with the Nanozoomer 2.0RS Hamamatsu Photonics and NDP.scan software (Hamamatsu City, Japan).

### Immunofluorescence

Immunofluorescent staining was performed on 8 µm tissue sections cryostained from OCT-frozen blocks. The slides were fixed in cold acetone (-20°C) for 20 minutes before a saturation step in PBS 3% BSA for 45 minutes. The labeling mix contained B220-FITC (REA755, Miltenyi Biotec®, Bergisch Gladbach, Germany) and CD3-PE (REA641, Miltenyi Biotec®, Bergisch Gladbach, Germany) antibodies diluted 1:100 and 1:167 respectively, and was incubated overnight at 4°C in the dark. The following day, DAPI staining was performed BD Biosciences®, Franklin Lakes, NJ). Slides were mounted and scanned using an epifluorscence microscope Nikon NiE (Nikon®, Tokyo, Japan). Labels were analyzed and quantified using QPath software.

### Cytokines production

Cytokine secretion levels were analyzed by flow cytometry. 2 million cells were cultured in complete RPMI medium (Eurobio Scientific, Essonnes, France) supplemented with 10% of FBS, 2 mM of L-Glutamine, 1% of Na pyruvate, 100 U/ml of penicillin and 100 μg/ml of streptomycin (ThermoFisher Scientific Waltham, MA) in 24-well plates. In each well, 4 µL of cocktail containing PMA/ionomycin + Brefeldin A (Biolegend, San Diego, CA) was added, as well as 2 µL of Monensin (Biolegend, San Diego, CA). Cells were cultured for 6 hours at 37°C, 5% CO2. Cells were then harvested and stained with surface CD19, CD3, CD4, CD8, and CD25 antibodies, followed by fixation and permeabilization using a commercial cytofix/cytoperm kit (BD Biosciences, USA) at 4°C for 30 min. Subsequently, intracellular FoxP3, IL-2, IFNγ, Granzyme B, IL-10, IL-6 and TNFα were stained at 4°C for another 30 min before proceeding to flow cytometry. Cells with no stimulation were set as negative controls (Supplementary Materials and Methods).

### Statistical analysis

Mann Whitney two-tailed tests were used for statistical analysis using GraphPad Prism software (*p<0.05, **p<0.01, ***p<0.001).

## Results

### The global proportion of immune B and T compartments was disturbed in *Myd88^L252P^* mice

From 6 months of age, blood samples were taken, every month, from the animals to monitor the presence of an Ig peak in the serum using serum protein electrophoresis. When animals show a monoclonal Ig peak in serum (10-12 months of age), they were euthanized for analysis. To study the tumour immune microenvironment and particularly the T-cell compartment, we first performed immunochemistry (IHC) staining with a CD3 antibody. In comparison with normal spleen from *Cd19^Cre^* control mice, we observed a reduction of CD3^+^ cells in *Myd88^L252P^* mice (Figure 1A). By using flow cytometry, we then evaluated the proportions of lymphoid cells in the spleens of *Cd19^Cre^* control and *Myd88^L252P^* mice. We confirmed an increase in the percentage of CD19^+^ B cells in tumours, representing 90% of spleen lymphocytes versus 60% under normal conditions. This was associated with a high decrease in CD4^+^ and CD8^+^ T cells (Figure 1B). These first results show that the B specific expression of *Myd88* mutation disturbed the immune compartment in spleen.

**Figure 1:**
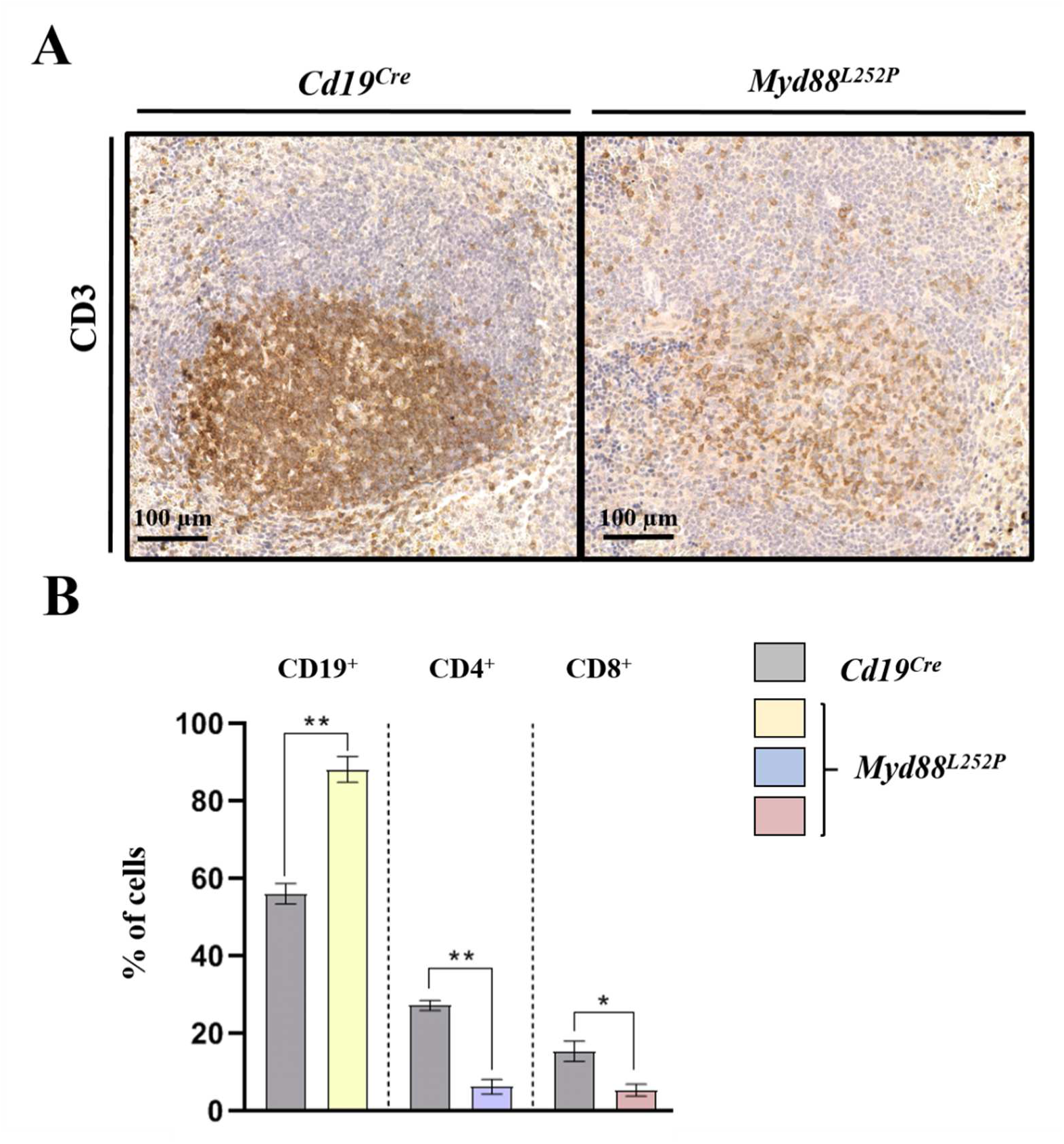
Splenic lymphocytes homeostasis from *Myd88^L252P^* mice was disrupted. Figure 1A: Anti-CD3 labelling was realised on paraffin-embed spleen sections and compared between *Cd19^Cre^* and *Myd88^L252P^* mice. Figure 1B: Percentages of CD19^+^, CD4^+^ and CD8^+^ cells among total splenic lymphocytes. *Cd19^Cre^* n=5; *Myd88^L252P^* n=5. Mann Whitney test p-value < 0.05 and p-value < 0.01 are symbolized by * and ** respectively.

### Tumour B cells displayed the hallmarks of an immunomodulatory phenotype

Analysis of *Myd88^L252P^* tumour B cells showed abnormal activity characterised by the overexpression of the activation markers CD80 and CD86 (Figure 2A-B). In addition, we analysed by flow cytometry the expression of two molecules strongly involved in the control of the anti-tumour immune response: the well-known immune checkpoint PD-L1; and the MHC class II complex, which enables antigen presentation to T cells. We then observed that tumour B cells overexpressed PD-L1, with 40% of them expressing it on average, compared with 10% for normal B cells (Figure 2C). In addition, B cells expressing the *Myd88^L252P^* mutation partially lost MHC class II expression (Figure 2D). Finally, we also characterized the cytokine secretion of these cells by flow cytometry. Interestingly, we found increased secretion of immunosuppressive cytokines like IL-10, IL-6 and TNFα, clearly demonstrating that splenic B cells of *Myd88^L252P^* mice exhibited an immunosuppressive phenotype (Figure 2E-H).

**Figure 2:**
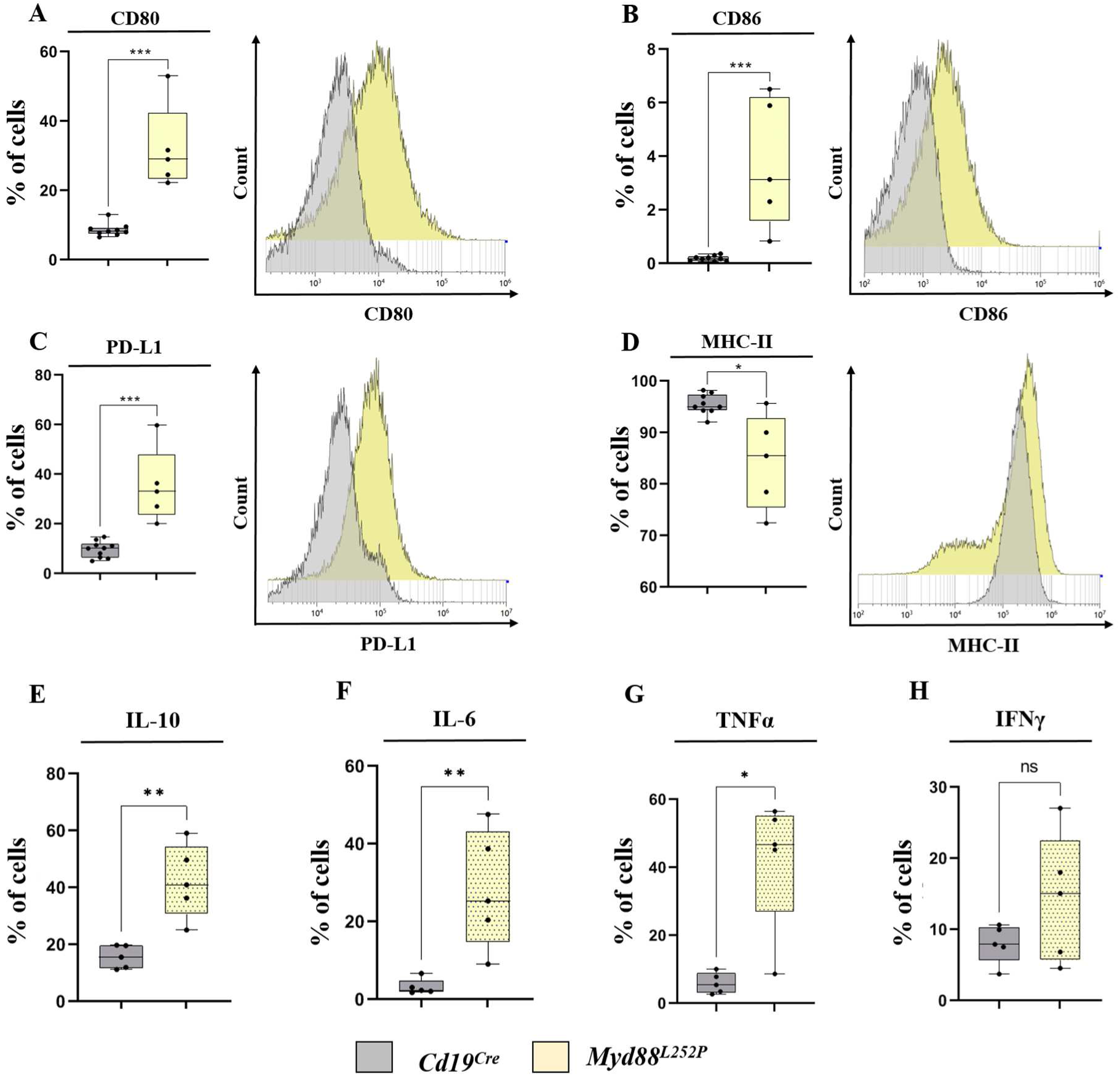
Tumor B cells from *Myd88^L252P^* mice presented an immunomodulatory phenotype. Figure 2A: Percentage of CD80 positive cells among CD19^+^ splenic population is shown on left panel. On the right panel, example of a single-parameter histogram showing the mean fluorescence intensity of the CD80 marker for *Cd19^Cre^* and *Myd88^L252P^* groups. Figure 2B: Percentage of CD86 positive cells among CD19^+^ splenic population is shown on left panel. On the right panel, example of a single-parameter histogram showing the mean fluorescence intensity of the CD86 marker for *Cd19^Cre^* and *Myd88^L252P^* groups. Figure 2C: Percentage of PD-L1 positive cells among B220^+^ splenic population is shown on left panel. On the right panel, example of a single-parameter histogram showing the mean fluorescence intensity of the PD-L1 marker for *Cd19^Cre^* and *Myd88^L252P^* groups. Figure 2D: Percentage of MHC-II positive cells among B220^+^ splenic population is shown on left panel. On the right panel, example of a single-parameter histogram showing the mean fluorescence intensity of the MHC-II marker for *Cd19^Cre^* and *Myd88^L252P^* groups. Figure 2E: Percentage of IL-10 positive cells among CD19^+^ splenic population. Figure 2F: Percentage of CD80 positive cells among CD19^+^ splenic population. Figure 2G: Percentage of TNFα positive cells among CD19^+^ splenic population. Figure 2H: Percentage of IFNγ positive cells among CD19^+^ splenic population. For all panels *Cd19^Cre^* n= 5 or 10; *Myd88^L252P^* n=5. Mann Whitney test p-value < 0.05, p-value < 0.01 and < 0.001 are symbolized by *, ** and *** respectively.

### An anti-tumour immune response was induced with signs of T cell activation

The immunosuppressive phenotype of tumour B cells should disrupt the anti-tumour immune response by affecting their recognition by T cells. Therefore, we continued the immune microenvironment study by looking at the localization of lymphoid compartments in the spleen comparing tumours from *Myd88^L252P^* mice with normal spleens. In contrast to our previous IHC analysis, immunofluorescence staining allowed us to distinguish these two populations using B220 and CD3 surface markers in the same spleen section. We then observed, in *Cd19^Cre^* control mice, a very close but distinct spatial distribution of B and T cells (Figure 3A-D). Indeed, B cells were organized in follicles, with T cells on the outside (Figure 3B-C). However, in tumour spleens of *Myd88^L252P^* mice, localization of B and T cells was disorganized with a mix of these two populations, suggesting migration of T cells towards the tumour area (Figure 3E-H). The presence of both CD4 and CD8-expressing T cells suggests that immune phenotype of cancer is an inflamed phenotype characterized by a pre-existing anti-tumour response that has been arrested by immunosuppressive processes.

**Figure 3:**
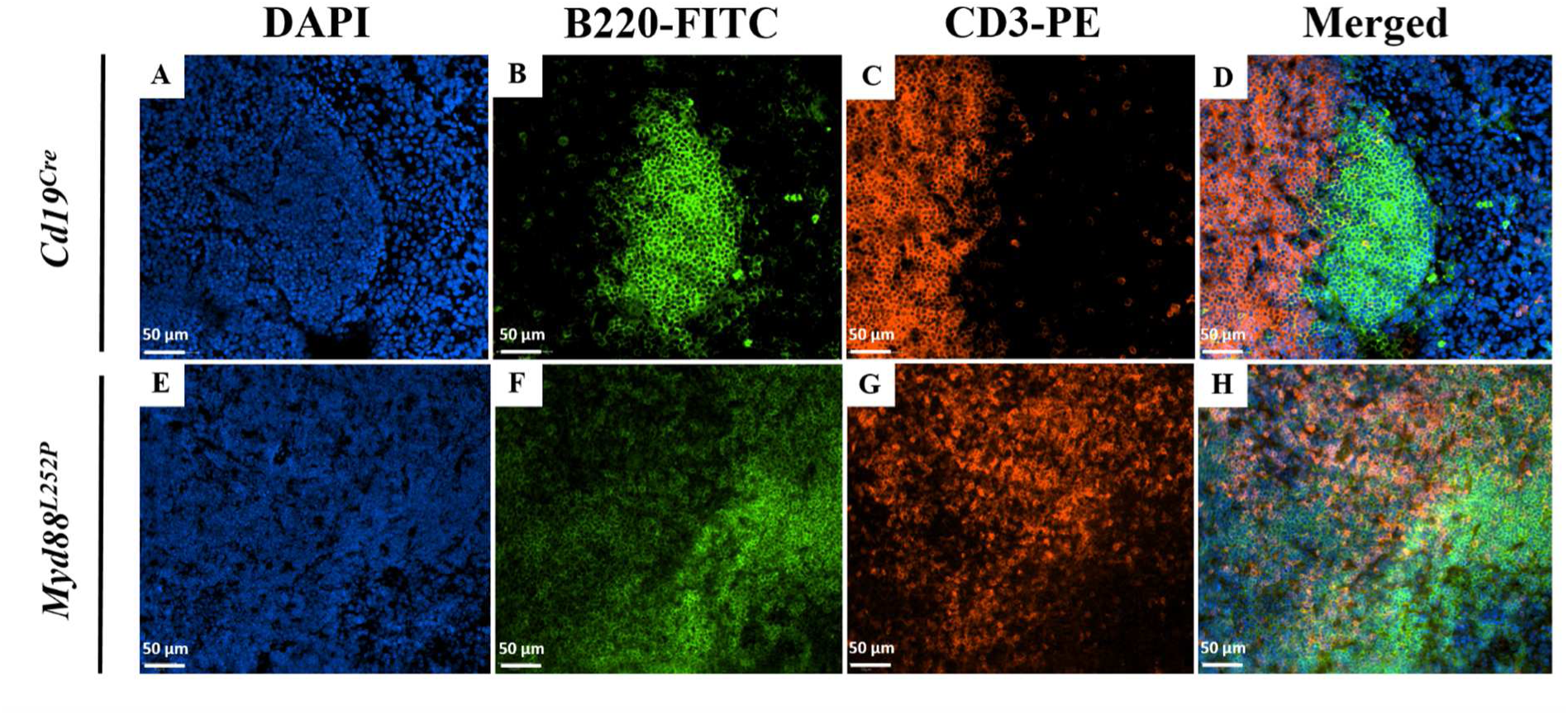
Immunofluorescence staining showed a dispruption of spleen organisation in *Myd88^L252P^* mice. Figure 3A and E: DAPI counter stain of *Cd19^Cre^* and *Myd88^L252P^* spleens respectively. Figure 3B and F: B cells of *Cd19^Cre^* and *Myd88^L252P^* mice are reveled with anti-B220-FITC antibody. Figure 3C and G: T cells of *Cd19^Cre^* and *Myd88^L252P^* mice are reveled with anti-CD3-PE antibody. Figure 3D and H: Visualisation of merged chanels (DAPI + FITC + PE).

Given the close proximity between T cells and tumour B cells, CD4 and CD8 populations were further characterized by flow cytometry. Double CD44-CD62L staining performed on CD4 and CD8 T cells distinguished three subpopulations: naive T cells (CD62L^+^ CD44^-^), memory T cells (CD62L^+^ CD44^+^) and effector T cells (CD62L^-^ CD44^+^). In *Myd88^L252P^* mice, both CD4^+^ and CD8^+^ populations showed a decrease in naive and memory compartments, with a high increase in effectors (Figure 4A-B). This increase was associated with activation of both CD4^+^ and CD8^+^ populations, revealed by increased expression of CD134 and CD137 markers (Figure 4C). We then investigated the activity of T cells at the functional level. *Ex vivo* experiments for the measurement of cytokine production were performed on splenocytes from *Cd19^Cre^* and *Myd88^L252P^* mice. We first focused our analysis on cytokines known responsible for the T-cell activation. Regarding the CD4-expressing T cells, no difference was observed in IL-2 secretion between *Cd19^Cre^* and *Myd88^L252P^* mice (Figure 5A). However, we detected an increase in IFNγ secretion in *Myd88^L252P^* mice (Figure 5B). CD8-expressing T cells, on the other hand, expressed significant amounts of both IL-2 and IFNγ (Figure 5C-D). We also found a higher secretion of granzyme B-type cytotoxic molecule (Figure 5E). In conclusion, these results show that tumour infiltrated T cells were mainly activated effector T cells with some features of Th1 polarization (high secretion of IL-2 and INF-γ), which confirm an engagement in an anti-tumour immune response with secretion of activating and cytotoxic cytokines.

**Figure 4:**
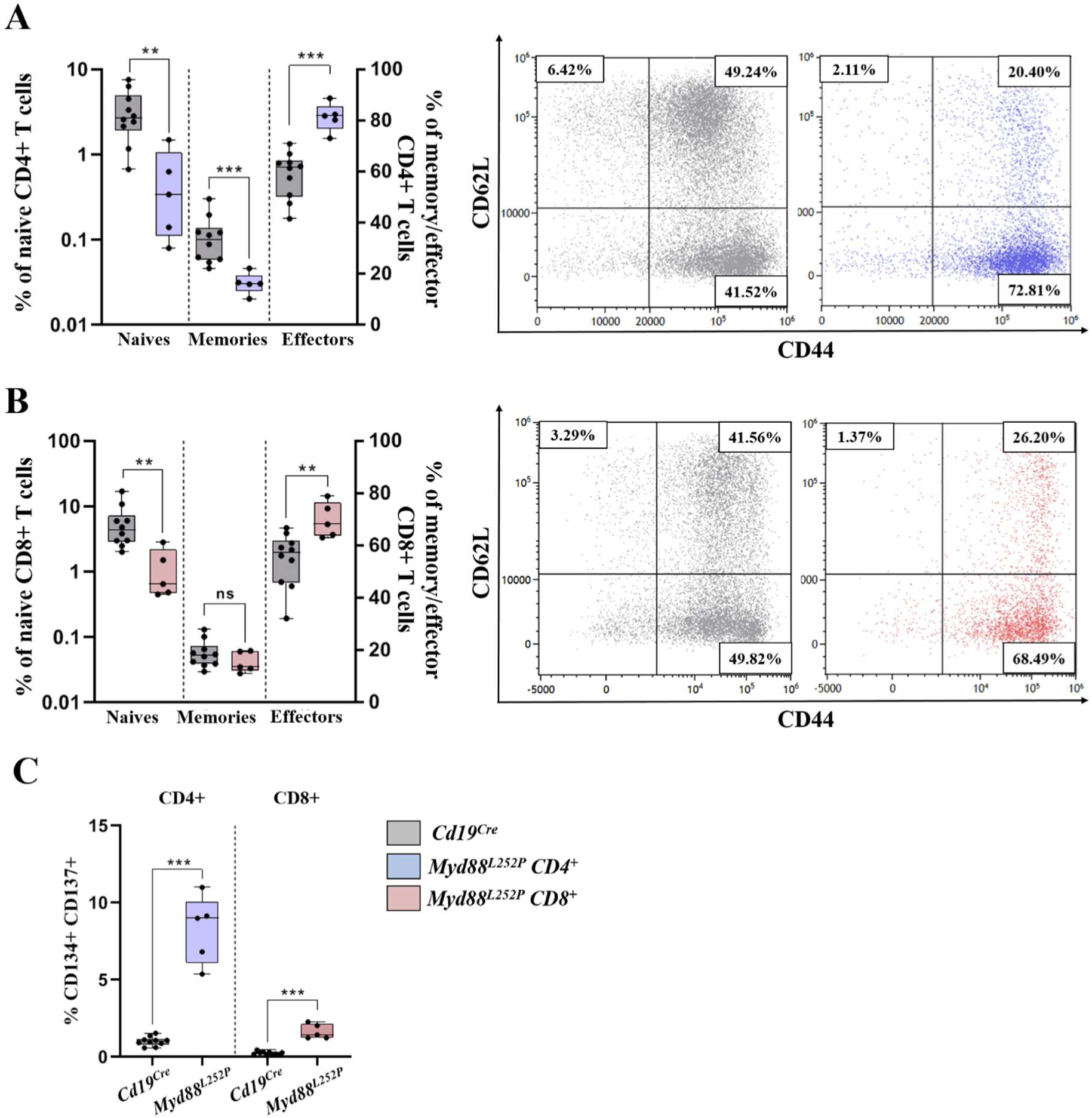
Tumor infiltrated T cells from *Myd88^L252P^* tumors presented an over-activated phenotype. Figure 4A: Left panel, percentages of naive (CD62L^+^ CD44^-^), memory (CD62L^+^ CD44^+^) and effector (CD62L^-^ CD44^+^) CD4 T cells are compared between *Cd19^Cre^* and *Myd88^L252P^* mice. Right panel, example of bi parametric flow cytometry histograms gated on CD4 positive T cells for expression of CD62L and CD44 for *Cd19^Cre^* and *Myd88^L252P^* mice. Figure 4B: Left panel, percentages of naive (CD62L^+^ CD44^-^), memory (CD62L^+^ CD44^+^) and effector (CD62L^-^ CD44^+^) CD8 T cells are compared between *Cd19^Cre^* and *Myd88^L252P^* mice. Right panel, example of bi parametric flow cytometry histograms gated on CD8 positive T cells for expression of CD62L and CD44 for *Cd19^Cre^* and *Myd88^L252P^* mice. Figure 4C: Percentages of CD4 and CD8 T cells expression both activation markers CD134 and CD137 are compared between *Cd19^Cre^* and *Myd88^L252P^* groups. For all panels, *Cd19^Cre^* n=10; *Myd88^L252P^* n=5. Mann Whitney p-value < 0.01 and < 0.001 are symbolized by ** and *** respectively.

**Figure 5:**
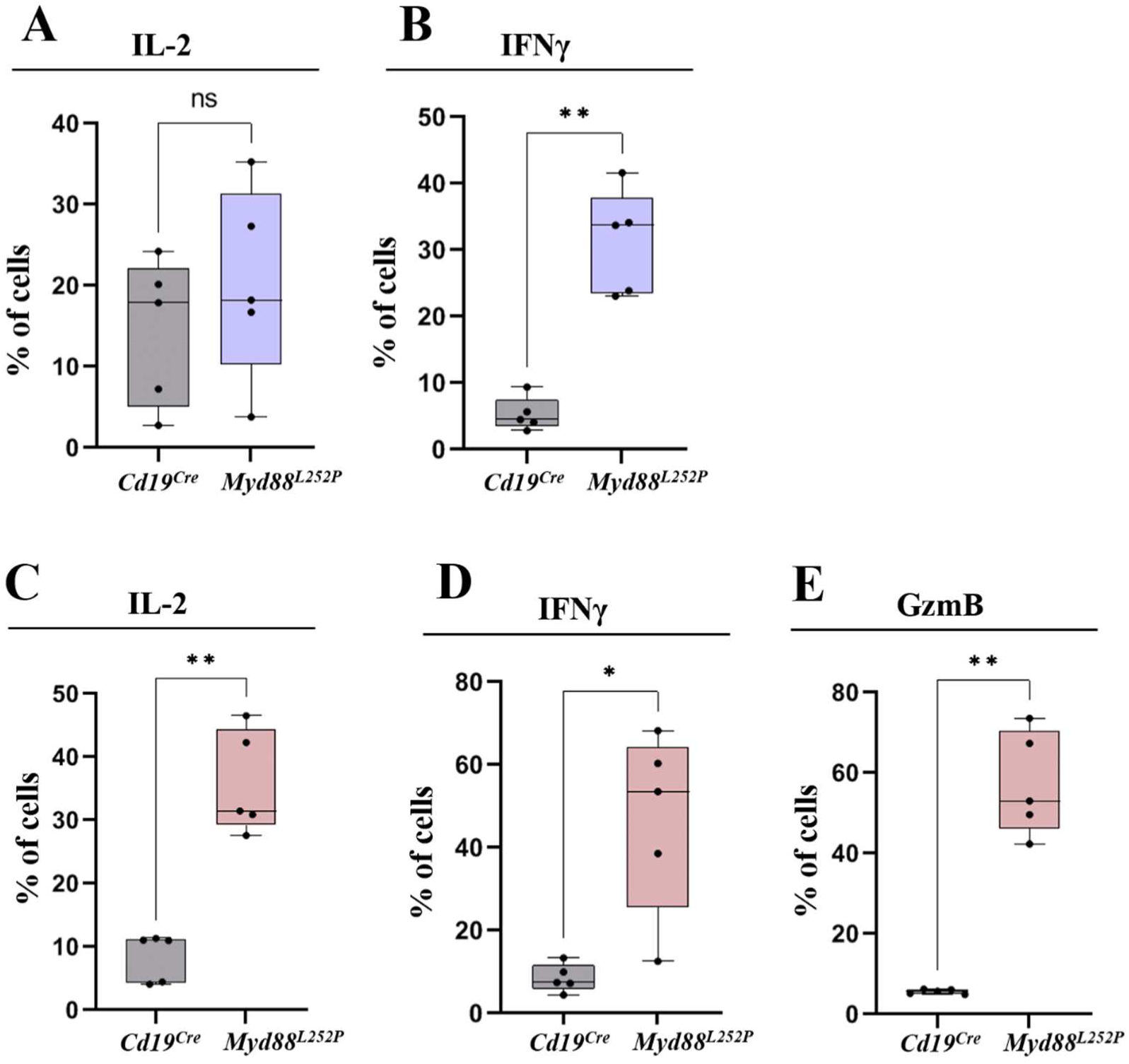
Over-activated T cells from *Myd88^L252P^* tumors presented a Th1 polarization. Figure 5A: Percentage of IL-2 positive CD4 T cells compared between *Cd19^Cre^* and *Myd88^L52P^* groups. Figure 5B: Percentage of IFNγ positives CD4 T cells compared between *Cd19^Cre^* and *Myd88^L52P^* groups. Figure 5C: Percentage of Granzyme B positives CD8 T cells compared between *Cd19^Cre^* and *Myd88^L52P^* groups. Figure 5D: Percentage of IL-2 positives CD8 T cells compared between *Cd19^Cre^* and *Myd88^L52P^* groups. Figure 5E: Percentage of IFNγ CD8 T cells compared between *Cd19^Cre^* and *Myd88^L52P^* groups. For all panels, *Cd19^Cre^* n=10; *Myd88^L252P^* n=5. Mann Whitney test p-value < 0.05 and p-value < 0.01 are symbolized by * and ** respectively.

### Tumour-infiltrated T cells were exhausted in *Myd88^L252P^* mice

The immunomodulatory phenotype of B cells and the involvement of T cells in an anti-tumour immune response suggested that the tumour must have developed mechanisms to limit T cell activity. To identify them, we better characterized T cells by studying surface expression of the four main checkpoints molecules that reflected T-cell exhaustion: Programmed cell Death-1 (PD-1), Cytotoxic T-Lymphocyte-Associated protein-4 (CTLA-4), T-cell Immunoglobulin and Mucin containing protein-3 (TIM-3) and Lymphocyte-Activation Gene-3 (LAG-3). Since B cells strongly expressed the PDL-1 molecule, we first investigated PD1 expression on CD4 and CD8-expressing T cells. We found a clear overexpression of PD-1 on the surface of both T-cell populations, with around 90% of CD4^+^ or CD8^+^ T cells expressing it in *Myd88^L252P^* mice, compared with 15% and 5%, respectively for CD4^+^ and CD8^+^ cells, in *Cd19^Cre^* control mice (Figure 6A). We completed this analysis by exploring other known markers of exhaustion. CTLA-4 expression was more heterogeneous, but overexpression can be found on both CD4^+^ and CD8^+^ cells (Figure 6B). Finally, TIM-3 and LAG-3 were expressed in far greater numbers by CD4 and CD8-expressing T cells in spleen of *Myd88^L252P^* mice than in *Cd19^Cre^* healthy tissue, where they are almost non-existent (Figure 6C-D). Moreover, analysis of cytokine secretion revealed overexpression of TNFα and IL-10 by both CD4^+^ and CD8^+^ T cells in *Myd88^L252P^* mice (Figure 6E-F). These results demonstrate the presence of CD4 and CD8 T-cell exhaustion in our mouse model. This exhaustion impaired the anti-tumour immune response, and appeared to be partly caused by the tumour cells themselves, which displayed an immunosuppressive phenotype.

**Figure 6:**
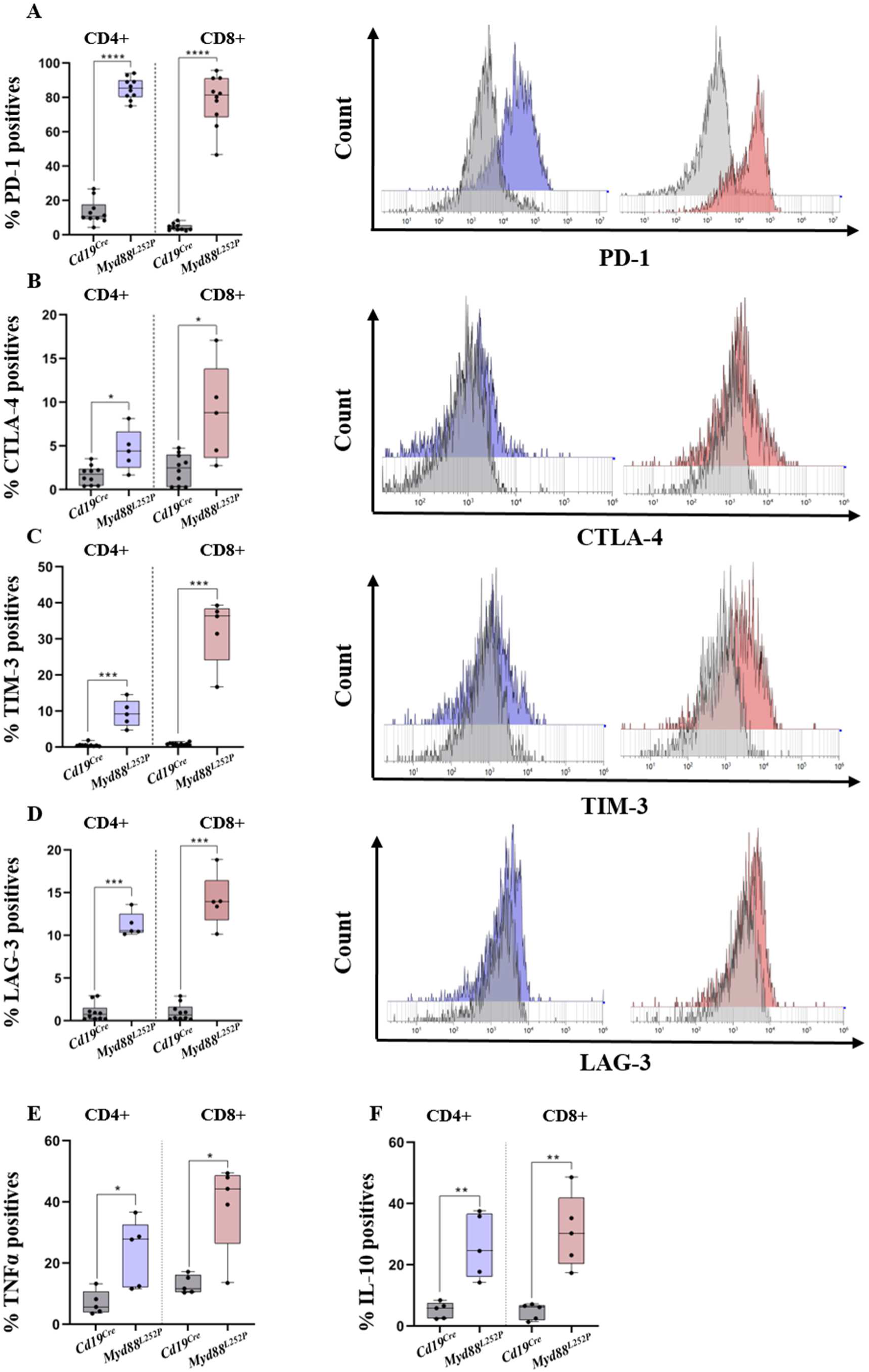
Infiltrated T cells from *Myd88^L252P^* tumors were exhausted. Figure 6A: Percentage of CD4 and CD8 T cells expressing PD-1 compared between *Cd19^Cre^* and *Myd88^L252P^* groups. Comparison of PD-1 MFI is shown on the right side of this panel. Figure 6B: Percentage of CD4 and CD8 T cells expressing CTLA-4 compared between *Cd19^Cre^* and *Myd88^L252P^* groups. Comparison of CTLA-4 MFI is shown on the right side of this panel. Figure 6C: Percentage of CD4 and CD8 T cells expressing TIM-3 compared between *Cd19^Cre^* and *Myd88^L252P^* groups. Comparison of TIM-3 MFI is shown on the right side of this panel. Figure 6D: Percentage of CD4 and CD8 T cells expressing LAG-3 compared between *Cd19^Cre^* and *Myd88^L252P^* groups. Comparison of LAG-3 MFI is shown on the right side of this panel. Figure 6E and F: Percentages of CD4 and CD8 T cells secreting TNFα and IL-10 compared between *Cd19^Cre^* and *Myd88^L252P^*. For all panels *Cd19^Cre^* n = 10; *Myd88^L252P^* n = 5. Mann Whitney test p-value < 0.05, p-value < 0.01 and < 0.001 are symbolized by *, ** and *** respectively.

### Increase of regulatory T cell population in *Myd88^L252P^* mice

Being known to be potent mediators of the inhibition of T-cell immune response, we finally focused on regulatory Tregs in our mouse model. Murine Tregs were defined by the simultaneous expression of CD4, CD25 and FoxP3 markers. Triple-positive cells were therefore analysed by flow cytometry. As a result, we found a significantly increased proportion of Tregs in *Myd88^L252P^* mice with 25% of Tregs among CD4^+^ versus 10% in healthy spleens (Figure 7A). The primary role of Tregs is to inhibit the T-cell immune response, notably by secreting anti-inflammatory cytokines. Thus, while Tregs normally express IL-6 and IL-10, we demonstrated that in the *Myd88^L252^* mice, Tregs overexpressed these two cytokines (Figure 7B-C). Finally, our results show that the percentage of Tregs expressing the TIM-3 marker was increased by a factor of 5 to 6 in tumours, compared with normal splenic tissue (Figure 7D). Moreover, TIM-3-positive Tregs secrete more IL-6 and IL-10 than TIM-3-negative Tregs (Figure 7E-F) revealing an increase immunomodulatory activity compared to TIM-3 negative Treg. In conclusion, these data have highlight an additional mechanism of immune escape through the expansion of the Treg population expressing the TIM-3 molecule.

**Figure 7:**
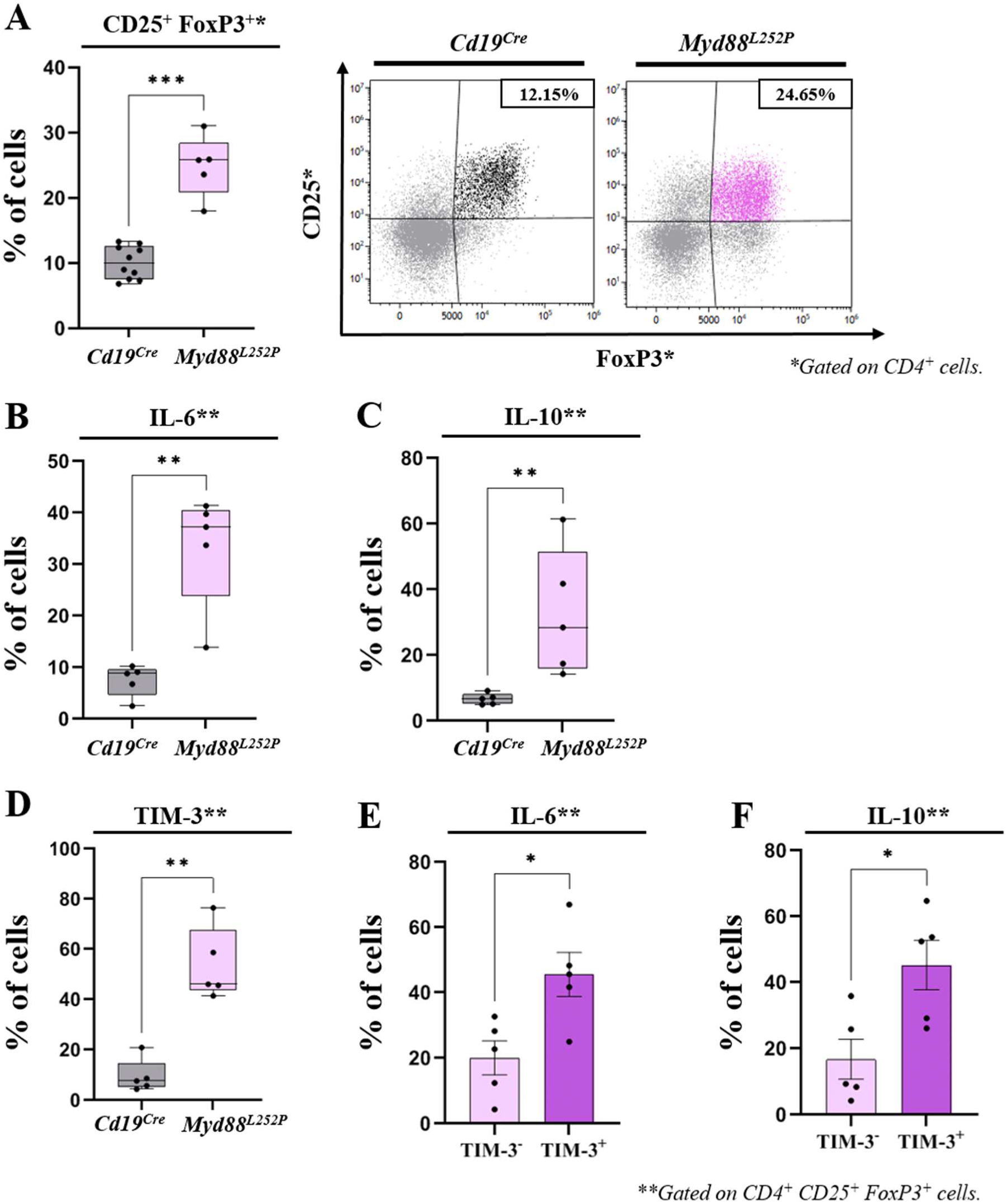
abnormal proportion of Tregs associated to a dysregulated anti-inflammatory phenotype support *Myd88^L252P^* tumor progression. Figure 7A: Left panel, percentage of CD25^+^ FoxP3^+^ Tregs among CD4^+^ T cells compared between *Cd19^Cre^* and *Myd88^L252P^* groups. Right panel, example of bi parametric flow cytometry histograms gated on CD4 positive T cells for expression of CD25 and FoxP3 for *Cd19^Cre^* and *Myd88^L252P^* mice. Figure 7B: Percentage of IL-6 secreting Tregs compared between *Cd19^Cre^* and *Myd88^L252P^* groups. Figure 7C: Percentage of IL-10 secreting Tregs compared between *Cd19^Cre^* and *Myd88^L252P^* groups. Figure 7D: Percentage of Tregs expressing TIM-3 compared between *Cd19^Cre^* and *Myd88^L252P^* groups. Figure 7E: Percentage of IL-6 secreting cells compared between TIM-3 positives and TIM-3 negatives Tregs. Figure 7F: Percentage of IL-10 secreting cells compared between TIM-3 positives and TIM-3 negatives Tregs. For all panels *Cd19^Cre^* n=5; *Myd88^L252P^* n=5. Mann Whitney test p-value < 0.05, p-value < 0.01 and < 0.001 are symbolized by *, ** and *** respectively.

## Discussion

The pressure of the immune surveillance is certainly one of the major driving forces in the emergence of B-cell lymphomas. Tumour cells are able to evade immune response by developing an immunosuppressive microenvironment through the dysregulation of immune checkpoint proteins expression, such proteins being essential in the negative control of activated immune cells. In our *Myd88^L252P^* mouse model that developed WM-like lymphoplasmacytic lymphoma, the TME study highlights the development of several immune escape mechanisms. First, tumour B cells displayed the hallmarks of immunosuppressive cells. As previously described, the B-cell expression of the *Myd88^L252P^* mutation led to the expansion of activated B cells which highly expressed the CD80 and CD86 markers; and this was associated with the overexpression of the PD-L1 molecule and the loss of MHC-II expression, sign of some loss of tumour-cell immunogenicity. We also confirm this immunosuppressive character by showing that tumour cells secreted large amounts of the immunosuppressive cytokines IL-10, IL-6 and TNFα. These results were consistent with the constitutive activation of several pathways such as NF-κB and JAK/STAT in our mouse model due to the expression of the *Myd88^L252P^* mutation. Moreover, these first results suggest that anti-tumour immune response should be disturbed and in particularly the T-cell functions. We actually detect some alterations in T-cell organisation and activity. First, we detected a migration of T cells close to the tumour B cells, suggesting the presence of a pre-existing anti-tumour immune response that was arrested, the exact definition of the immune-inflamed phenotype reviewed by Chen and Mellman [26]. In a second step, we characterized those T cells in details and we found that both CD4 and CD8 expressing T cells were mainly effector T cells with a loss in T-cell memory compartment. Moreover, those T cells showed all the signs of a Th1-type anti-tumour immune response with IL2, IFNγ and TNFα secretion. In parallel, we also demonstrated the existence of a T-cell exhaustion with an overexpression of the four main immune checkpoint molecules PD-1, CTLA-4, TIM-3 and LAG-3, associated with an increase in TNFα and d’IL-10 secretion. In our mouse model, we confirmed what was suspected in WM patients: the presence of the constitutively active mutated MYD88 protein seems to induce the emergence of an immunosuppressive tumour phenotype responsible for T-cell exhaustion with age [25], [27]. Furthermore, it would be particularly interesting to determine exactly which antigens are recognized by the exhausted T cells in our model, as this would perhaps enable us to draw up a list of antigens favouring T-cell exhaustion and tumour escape. Searching for these antigens in the patient would provide a predictive element for T-cell exhaustion, and they could also constitute new therapeutic targets, since it is conceivable that blocking these antigens would prolong the efficacy of the anti-tumour immune response.

Moreover, similar to the observations made in WM patients, we demonstrated in our mouse model an increase in the population of Tregs in splenic tumours [25]. Tregs are immunosuppressive cells, and deregulation of their functions is known to be involved in many cancers. We showed here that Tregs from MW-like tumours overexpressed the anti-inflammatory cytokines IL-10 and IL-6. This is in agreement with a recent study showing, in another mouse model and in patient, the existence of a Treg-mediated immunosuppressive phenotype in WM [28]. We also discovered in our mouse model, in comparison to control mice, an expansion of TIM-3^+^ Tregs cells which secreted higher amounts of immunosuppressive IL-6 and IL-10 cytokines than do TIM-3^-^ Tregs. To note, TIM3 positive Tregs are shown to be associated with non-favourable prognostic in many cancers due to their huge secretion of IL-10 [29]. Targeting this population would then be a therapeutic prospect for WM. In this sense, Sacco et al. propose to target the CD40/CD40L axis after having identified its role in the WM cell-Tregs interaction [28].

All these results contribute to better understand tumour microenvironment in WM with the aim to identify novel mechanisms of B-cell immune escape, which may be the keystone of future therapies. Targeting immune checkpoints and especially PD-L1 gave spectacular results in various solid cancers such as those of lung or colon [30]–[32]. Apart Hodgkin lymphomas, such therapies did not reach the hopes in B-cell lymphomas, especially for those with NF-κB activation. This lack of curative effect in these patients, despite PD-L1 expression by tumour cells in most cases, is to date poorly understood. In WM, such therapies could be of great help because of the *MYD88* mutation that constitutively activates NF-κB. However, failures in other B-cell cancers prevent any attempts of such new therapeutic proposal until immune surveillance escape will be deciphered and understood. Mouse models are thus useful for the preclinical study of these therapies, evidencing the interest of combinations that include treatments able to restore the anti-tumour immune response. Before trying such strategies in Humans, this is testable in genetically engineered mouse models. Our *Myd88^L252P^* model would then be of interest and will be a very good pre-clinic tool for testing new therapeutic combinations combining current therapies with those targeting immune system reactivation, such as anti-PD-1 or anti-TIM-3 immunotherapies or those directly targeting the binding of immunosuppressive cytokines IL-6 or IL-10 to their receptors.

## Author Contributions

QL, OT, MC and LJ performed and analysed experiments. QL performed flow cytometry analysis. CV-F contributed to the experiments and analysed the results. JF and CV-F directed the study. QL and CV-F wrote the manuscript. All authors contributed to the article and approved the submitted version.

## Acknowledgments

We thank the animal, histology, cytometry and microscopy facilities of the technological platform of the University of Limoges BISCEm (Inserm 042, CNRS 2015, Hospital University Center of Limoges).

