## Supplementary materials and methods for "B-cell specific expression of the murine *Myd88^L252P^* mutation correlates with the establishment of an immunosuppressive microenvironnement in Waldenström macroglobulinemia lymphoplasmacytic lymphoma"

**Details of panels used for B and T-cell characterization by flow cytometry**

| Marker | Fluorochrome | Clone | Supplier |
|---|---|---|---|
| **Activated B cells panel** | | | |
| CD80 | BV421 | 16-10A1 | Biolegend® |
| CD86 | PE Vio770 | REA1190 | Miltenyi® |
| CD19 | BV510 | 1D3 | BD® |
| IgD | PerCP Cy5.5 | 11-26c.2A | BD® |
| IgM | PE | R6-60.2 | BD® |
| B220 | BV785 | RA3-6B2 | Biolegend® |
| Viakrome 808 | | / | Beckman Coulter® |
| **PD-L1/MHC-II panel** | | | |
| B220 | BV421 | RA3-6B2 | Biolegend® |
| PD-L1 | PE | 10F.9G2 | Biolegend® |
| CMH-II | Viogreen | REA813 | Miltenyi® |
| Viakrome 808 | | / | Beckman Coulter® |
| **Activated T cells panel** | | | |
| CD4/CD8 | PE Dazzle 594 | RM4.5/53-6.7 | Biolegend® |
| CD134 | APC | OX86 | Tonbo Bioscience® |
| CD137 | PE | 17B5 | Biolegend® |
| Viiakrome 808 | | / | Beckman Coulter® |
| **T cells subgroups panel** | | | |
| CD4 | PE Dazzle 594 | RM4.5 | Biolegend® |
| CD8 | APC | 53-6.7 | Biolegend® |
| CD44 | PE | IM7 | Biolegend® |
| CD62L | BV421 | MEL14 | Biolegend® |
| Viakrome 808 | | / | Beckman Coulter® |
| **Exhausted T cells panel** | | | |
| CD4 | Vioblue | REA604 | Miltenyi® |
| CD8 | APC Vio770 | REA601 | Miltenyi® |
| CD3 | PE Vio770 | REA641 | Miltenyi® |
| CD45 | PerCP Vio700 | REA737 | Miltenyi® |
| PD-1 | PE | REA802 | Miltenyi® |
| CTLA-4 | PE Vio615 | REA984 | Miltenyi® |
| TIM-3 | APC | REA602 | Miltenyi® |
| LAG-3 | BV711 | C9B7W | Biolegend® |
| Viakrome 808 | | / | Beckman Coulter® |

**Details of panels used for the study of cytokine production by flow cytometry**

| Marker | Fluorochrome | Clone | Supplier |
|---|---|---|---|
| **B cells cytokines panel** | | | |
| CD19 | BV510 | 1D3 | BD® |
| IL-10 | BV421 | JES5-16E3 | Biolegend® |
| TNFα | APC Cy7 | MP6-XT22 | Biolegend® |
| IL-6 | APC | REA1034 | Miltenyi® |
| IFNγ | BV711 | XMG1.2 | Biolegend® |
| **T cells cytokines panel** | | | |
| CD4 | PE Dazzle 594 | RM4.5 | Biolegend® |
| CD8 | APC | 53-6.7 | Biolegend® |
| Granzyme B | FITC | GB11 | Biolegend® |
| IL-2 | PE | JES6-5H4 | Biolegend® |
| TNFα | APC Cy7 | MP6-XT22 | Biolegend® |
| IL-10 | BV421 | JES5-16E3 | Biolegend® |
| IFNγ | BV711 | XMG1.2 | Biolegend® |
| **Tregs cytokines panel** | | | |
| CD4 | FITC | RM4.5 | Biolegend® |
| CD3 | PE Vio770 | REA641 | Miltenyi® |
| FoxP3 | PE | REA768 | Biolegend® |
| TIM-3 | BV711 | B8.2C12 | Biolegend® |
| IL-10 | BV421 | JES5-16E3 | Biolegend® |
| IL-6 | APC | REA1034 | Biolegend® |
